# *In Silico:* Mutagenicity and carcinogenicity prediction of Sugar substitutes

**DOI:** 10.1101/2021.01.28.428696

**Authors:** Charli Deepak Arulanandam, Venkatadri Babu, Yugendhar Soorni, R Prathiviraj

## Abstract

Sugar substitutes are mostly artificial, man-made industrial products used as additives in food and beverages. Most of these substances flow through the digestive tract and food chains to become emerging organic contaminants in various abiotic and biotic environmental media. Here, we predict the mutagenicity and carcinogenicity of commonly used sugar substitutes using the *in silico* based methods. The simplified molecular-input line-entry system (SMILES) of sugar substitutes were obtained from the PubChem database for the toxicity predictions. Here, sixteen sugar substitutes tested out of these four compounds Glucin (GLU), and 5-nitro-2-propoxyaniline (P-4000), SCL, Ace were predicted as mutagens by using *in silico* tools such as LAZAR, pKCSM, and Toxtree. Based on the predicted results GLU and P-4000 were predicted as carcinogenic sugar substitutes.

## 1. INTRODUCTION

Obesity has become a major challenge for chronic disease prevention and health across the globe (Sylvetsky et al., 2012; Hruby and Hu, 2015) and sugar substitutes are increasingly replacing natural sugars as sweeteners (Jiang et al., 2018). The consumption of sugar substitute containing beverages among U.S. children increased twice (Sylvetsky et al., 2013). Adverse human health effects from the application of artificial sweeteners (ASs) have been reported in several studies. Also unfavorable embryonic development was observed in women consuming artificially sweetened coffee (Setti et al., 2017). Since several sugar substitutes were shown to have adverse effects on public health (Ma et al., 2009; Steinert et al., 2011; Ford et al, 2011; Durán et al., 2018), that was also indicated by animal studies (Bellringer et al., 1995; Swithers et al., 2012; Jiang et al., 2018; Haighton et al., 2018), and *in vitro* studies (Pepino, 2015). But, no side effects observed during appetite profiling involving human subjects for ASs (Steinert et al., 2011). Sugar substitutes are also reported from diverse environmental samples (Kokotou et al., 2012). They are either discarded as unused products or with unused food or beverages, or pass the digestive tract unchanged, end up in sewage systems and subsequently in watersheds, and may penetrate down to the groundwater (Yang et al., 2018). By conventional wastewater treatment processes, some of the sugar substitutes are difficult to degrade (Lin et al., 2017). Due to the high costs for material and time of experimental testing, it became convenient to develop robust based on the *in silico* approaches to perform chemical toxicity predictions such as the mutagenicity of compounds (Xu et al., 2012). Such computational methods are gaining increasing acceptance in toxicological risk assessments (Staal et al., 2017). As for toxicity evaluations, *in silico* studies are more fast and cost-efficient methods and have a unique advantage in being able to estimate the direct or side-effect toxicity (Madan et al., 2013). The present contribution aims to predict the mutagenic and carcinogenic properties of sugar substitutes by using the LAZAR, Toxtree, and CarcinoPred-EL.

## 2. MATERIALS AND METHODS

### 2.1. SMILES based input format for compounds

Molecular structures were represented by the strings of special characters using SMILES as kernel strings. Since SMILES-based similarity functions are computationally more efficient, we used a SMILES-based toxicity prediction (Öztürk et al., 2016; Frenzel et al., 2017). Sugar substitutes data were retrieved from the PubChem data bank for *in silico* assessments. SMILES of the sugar substitutes were provided in **Table 1**. Molecular structures represented in **Figure. 1** were made by using the Marvin Sketch structure drawing tool (www.chemaxon.com).

**Table 1.**
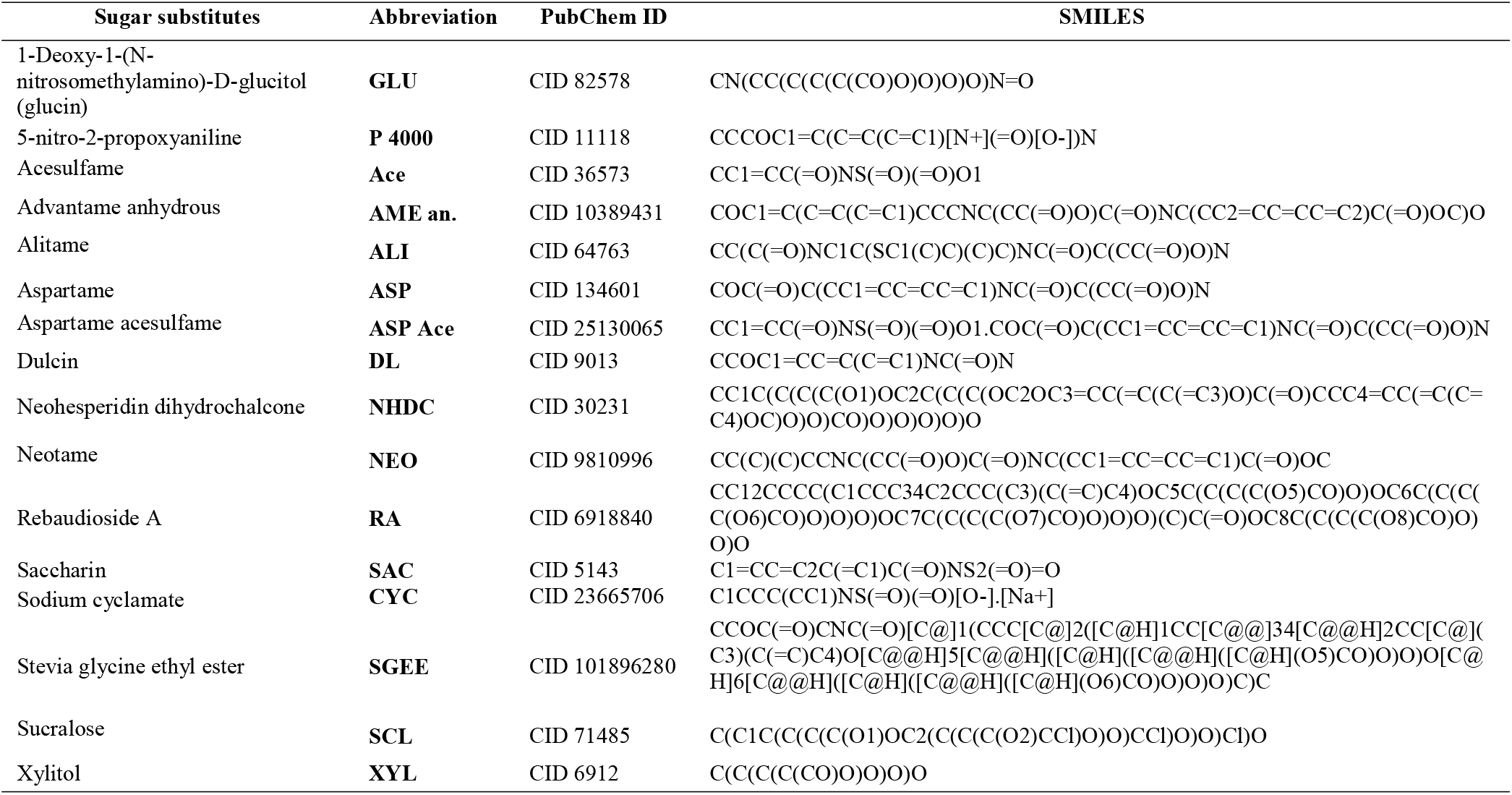
Sugar substitutes SMILES and PubChem ID.

**Figure. 1.**
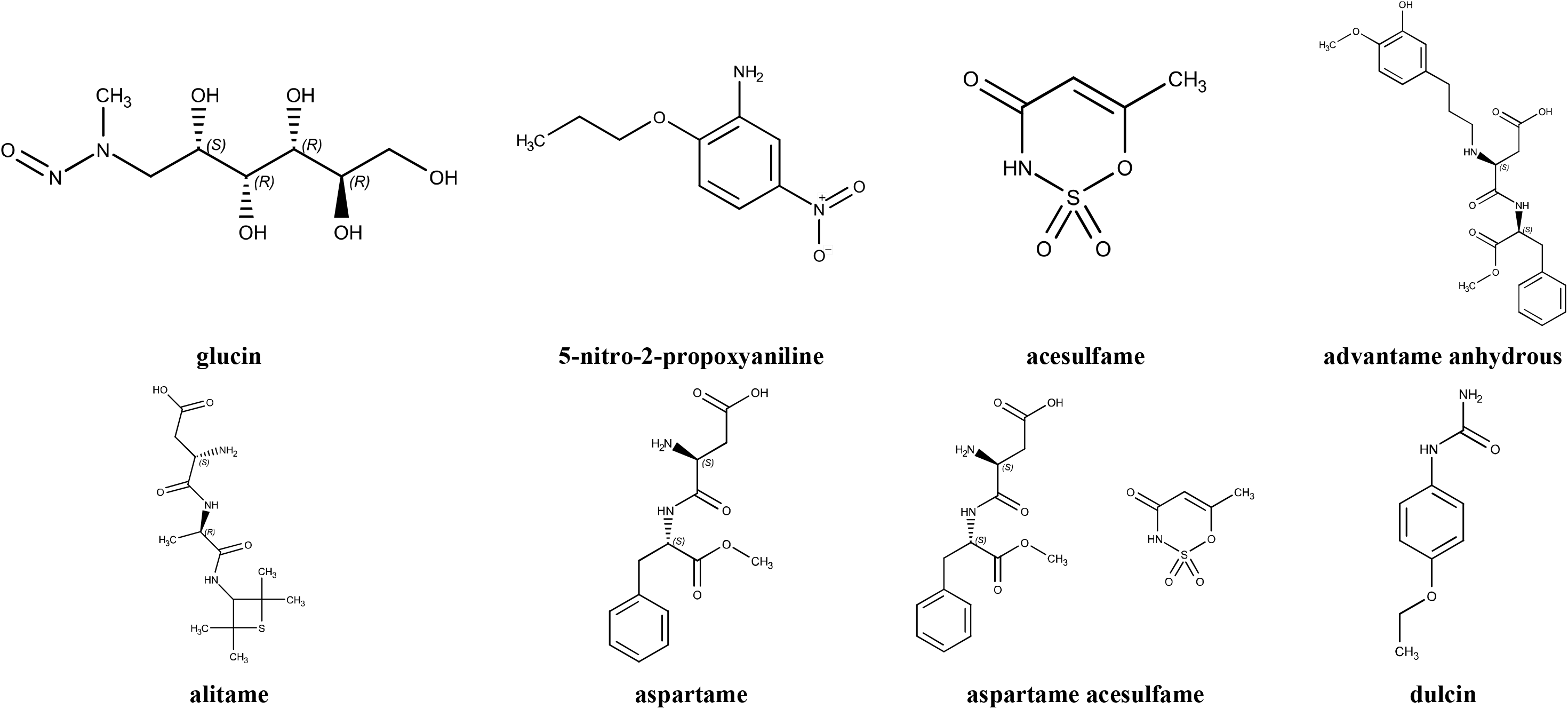
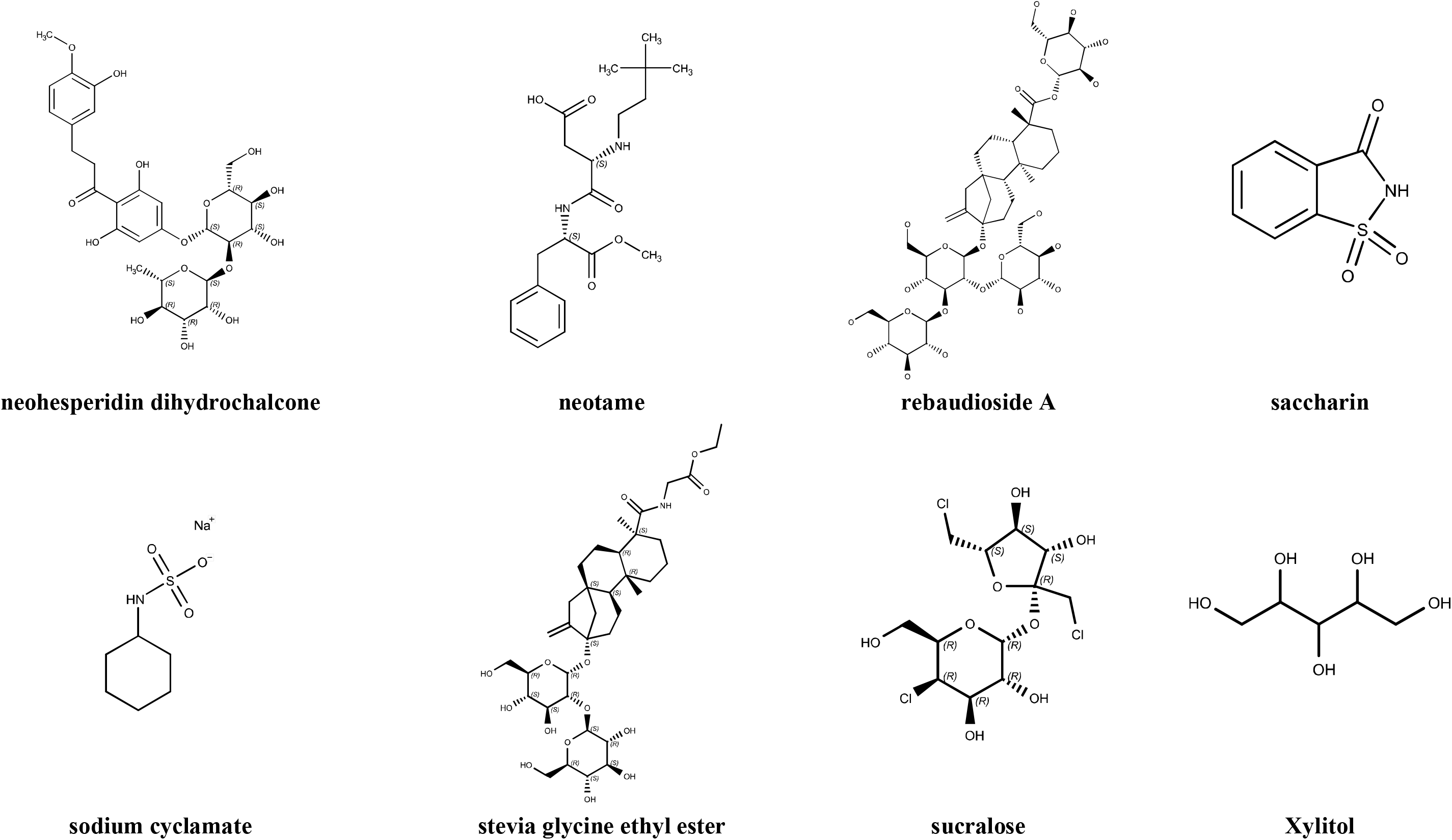
Structures of Sugar substitutes. Molecular structures of the sugar substitutes were drawn by MarvinSketch.

### 2.2. Mutagenicity and carcinogenicity prediction

In the toxicological endpoints of chemical substances, mutagenicity and carcinogenicity are of great concern because of their serious effects on human health. Mutagenicity represents one of the most important endpoints of toxicity and also indicates the permanent changes in the DNA sequence of an organism, which may result in heritable changes in the characteristics of organisms (Słoczyńska et al., 2014). The mechanism of action of mutagenic chemicals is mainly through DNA damage, which causes mutagenic effects such as chromosomal aberrations, frameshift mutation, and point mutations. Mutagens are not only involved in carcinogenesis and genotoxicity. They are also involved in the inception and pathogenesis of several chronic diseases including neurodegenerative disorders, diabetes, aging processes, arthritis, cardiovascular impairment, chronic inflammation, and hepatic disorders (Bhattacharya, 2011).

#### 2.3.1. LAZAR

LAZAR is a web server for the prediction of toxicity of known and unknown chemical structures (Helma, 2006). This web server provides a modular program for toxicity prediction (Maunz et al., 2013). LAZAR can predict the complex toxicological properties of compounds like mutagenicity, carcinogenicity, acute toxicity, maximum recommended daily dose, and blood-brain barrier penetration of chemical substances by using the data mining algorithms. This provides LAZAR with the advantage of providing toxicological predictions for endpoints that provide sufficient empirical information. LAZAR approach prevents that incomplete or incorrect background information is affecting the model predictions (Maunz et al., 2013). This web server is available from https://nano-lazar.in-silico.ch/predict.

#### 2.3.2. pKCSM

This tool is used to predict the Absorption Distribution Metabolism Excretion and Toxicity (ADMET) of chemical compounds (Pires et al., 2015). The pkCSM web server available from http://biosig.unimelb.edu.au/pkcsm/ that allows rapid evaluation of pharmacokinetic and toxicity properties of chemical compounds.

#### 2.3.3. Toxtree

Toxic Hazard Estimation by decision tree approach (Toxtree v2.6.13) provides toxic effect predictions. Toxtree was developed by Idea Consult Ltd. (Sofia, Bulgaria) to provide the structure-activity-relationships (SAR) for fine-tuned toxicity estimates. To obtain toxicity characteristics, we submit the SMILES of sugar substitute in the Toxtree tool (see Benigni et al., 2008; Gumber et al., 2018). The Toxtree web server and downloadable version of Toxtree v2.6.13 is available from http://toxtree.sourceforge.net.

## 3. RESULTS

In this study LAZAR, pKCSM, Toxtree tools were used to predict the mutagenicity and carcinogenicity of sugar substitutes. LAZAR, pKCSM, and Toxtree were used to predict the mutagenicity of test compounds. The following sugar substitutes were not indicated as mutagenic in this assessment: CYC, SGEE, ASP, NEO, SAC, AME an., ALI, NHDC, RA, and XYL were no mutagens as per the pKCSM and Toxtree prediction (**Table 2**). From the pKCSM and Toxtree predictions were SCL, Ace, GLU, and P-4000 as mutagens in both of the *in silico* models. ASP, Ace, and DL were predicted as non-mutagens by LAZAR and pKCSM but similar results not observed in the Toxtree.

**Table 2.** Mutagenicity prediction by pKCSM and Toxtree tool.

| Sugar substitutes | LAZAR | pKCSM | Toxtree |
| --- | --- | --- | --- |
| GLU | M | Yes | Yes |
| P 4000 | M | Yes | Yes |
| Ace | M | Yes | Yes |
| AME an. | NM | No | No |
| ALI | NM | No | No |
| ASP | NM | No | No |
| ASP Ace | NM | No | Yes |
| DL | NM | No | Yes |
| NHDC | NM | No | No |
| NEO | NM | No | No |
| RA | NM | No | No |
| SAC | NM | No | No |
| CYC | NM | No | No |
| SGEE | NM | No | No |
| SCL | M | Yes | Yes |
| XYL | NM | No | No |

*M/Yes-mutagen, NM/No-non mutagen, \*could not predict.*

Accordingly, the LAZAR, Toxtree, and CarcinoPred-EL prediction showed that CYC, SGEE, SAC, AME an., NHDC, and RA are not bearing a risk of cancer. ASP is predicted as a carcinogen by LAZAR *in silico* rodent and mouse model but not by LAZAR *in silico* rat model, Toxtree and CarcinoPred-EL. SCL was predicted as a carcinogen in the LAZAR *in silico* mouse model. Toxtree also predicts SCL as a carcinogen; this is not indicated in the LAZAR *in silico* rat model and CarcinoPred-EL. Some of the sugar substitutes predicted as a carcinogen in only one prediction model such as ALI which was predicted as a carcinogen in LAZAR *in silico* rodent model.

ASP and Ace was predicted as carcinogenic by the Toxtree carcinogenicity prediction. DL was predicted as a carcinogen in all tested carcinogenicity prediction models except the LAZAR *in silico* rat model. The two sugar substitutes-GLU, and P-4000 are predicted as carcinogens in all tested *in silico* carcinogenicity prediction models (**Table 3**).

**Table 3.** Sugar substitutes carcinogenicity prediction.

| Sugar substitutes | LAZAR |  |  | Toxtree | CarcinoPred-EL |  |  |
| --- | --- | --- | --- | --- | --- | --- | --- |
|  | Rodent | Mouse | Rat |  | RF | SVM | XGBoost |
| GLU | C | C | C | C | C | C | C |
| P 4000 | C | C | NC | C | C | C | C |
| Ace | * | * | * | C | NC | NC | NC |
| AME an. | NC | NC | NC | NC | NC | NC | NC |
| ALI | C | NC | NC | NC | NC | NC | NC |
| ASP | C | C | NC | NC | NC | NC | NC |
| ASP Ace | NC | NC | NC | C | NC | NC | NC |
| DL | C | C | NC | C | C | C | C |
| NHDC | NC | NC | NC | NC | NC | NC | NC |
| NEO | NC | C | NC | NC | NC | NC | NC |
| RA | NC | NC | NC | NC | NC | NC | NC |
| SAC | NC | NC | NC | NC | NC | NC | NC |
| CYC | NC | NC | NC | NC | NC | NC | NC |
| SGEE | NC | NC | NC | NC | NC | NC | NC |
| SCL | C | C | NC | C | NC | NC | NC |
| XYL | C | C | NC | NO | * | * | * |

*C-carcinogen, NC-non carcinogen, \*could not predict.*

## 4. DISCUSSION

Sugar substitutes such as saccharin being the known artificial sweetener. Next to saccharin, cyclamate and aspartame are the most important artificial sweeteners which were discovered in quite remarkable ways. Chemists experienced the sweet taste by accident while smoking a cigarette (David, 2007). In spite of visually harmful structural features, saccharin (o-sulfobenzoic acid imide) provides uniquely low toxicity. Reasons for its low toxicity are not known as yet. We found that Saccharin and rebaudioside A were not mutagenic and carcinogens according to our *in silico* predictions. Saccharin and rebaudioside A are widely used as non-caloric sweeteners. However, a study on weanling rats with daily exposure to saccharin sodium and rebaudioside-A showed an increase of abnormal estrous cycles, higher numbers of ovarian cysts, elevated serum progesterone levels, and increased expression of steroidogenesis-related factors in the treatments (Jiang et al., 2018). Conversely, rebaudioside A treated groups showed decreased serum progesterone levels. The findings of Jiang and coworkers (2018) suggest that saccharin sodium exerts adverse biological effects on rodent ovaries, and that rebaudioside A is a potential steroidogenic disruptor in female rats. High intake of sweetened beverages is also associated with adverse health outcomes in animal rat models. Consumption of artificially sweetened coffee was negatively associated with rat embryo quality on days 2 and 3. Exposure to rebaudioside A remarkably decreased total cholesterol in female rats (Curry and Roberts, 2008).

According to our toxicity assessment, neither aspartame nor saccharin has mutagenic nor carcinogenic properties. Four compounds (SCL, Ace, GLU, and P-4000) were suggested as mutagenic using the three *in silico* mutagenicity predictions. GLU and P-4000 were predicted as carcinogens in all tested *in silico* carcinogenicity prediction models. Based on the predicted results from above *in silico* risk assessments, Glucin (GLU) and 5-nitro-2-propoxyaniline (P-4000) were predicted as carcinogenic without exception in all *in silico* carcinogenicity prediction models applied here.

## 5. Conclusions

The present SMILES based low-calorie sweetener toxicity prediction study assessed the mutagenicity, and carcinogenicity of sugar substitutes. Sugar substitutes are detectable in all spheres of the earth, but mostly affect organisms in aquatic systems, from where they can return to and affect human consumers by drinking water or consuming agricultural or aquacultural food products. As such they provide important issues for food and drink safety. For the risk assessment of sugar substitutes, we made toxicity predictions that were based on SMILES kernel strings of the sweetener compounds which were analysed through the computational tools LAZAR, pKCSM, and Toxtree. From 16 sugar substitutes tested here, four compounds (SCL, Ace, GLU, and P-4000) were confirmed as mutagens by using three mutagenicity prediction tools. GLU and P-4000 were predicted as carcinogens in all tested *in silico* models applied here. The results provided predictions on the impacts of sugar substitutes on the abiotic and biotic environment including humans. It considered toxicological endpoints and provided a fast, inexpensive, and heuristic approach.

## ACKNOWLEDGMENTS

Charli Deepak Arulanandam expressed his appreciation to Yugendhar Chowdhary, Onium Life Sciences Pvt. Ltd, Bangalore, India for the data verification.

## Competing Interests Statement

The authors have no competing interests associated with this manuscript.

## Funding

This research was supported by the Kaohsiung Medical University Fellowship.

## Supporting Information

Supporting information can be accessed through www.chemdata.org

## ABBREVIATIONS USED

SMILES: Simplified molecular-input line-entry system
LAZAR: lazy structure–activity relationships)
pKCSM: predicting small-molecule pharmacokinetic properties
DNA: Desoxyribonucleic acid
CYC: sodium cyclamate
SGEE: Stevia glycine ethyl ester
ASP: aspartame
SCL: sucralose
NEO: neotame
Ace: acesulfame
SAC: saccharin
AME an.: advantame anhydrous
ALI: alitame
ASP Ace: aspartame acesulfame
DL: dulcin
GLU: glucin
NHDC: neohesperidin dihydrochalcone
P-4000: 5-nitro-2-propoxyaniline
RA: rebaudioside A
XYL: Xylitol
ECB: European Chemicals Bureau
SAR: structure-activity relationships
ADMET: Absorption Distribution Metabolism Excretion and Toxicity

## Notes

http://www.chemdata.org

### Competing Interest Statement

The authors have declared no competing interest.

